# Neural tracking of surprisal and semantic distance in naturalistic movie viewing

**DOI:** 10.64898/2026.09.09.749230

**Authors:** Ryan M. O’Leary, Hailey C. Smith, Emily B. Myers, Jamie Reilly, Jonathan E. Peelle

**Author notes:** Please address correspondence to: Dr. Jonathan Peelle, Institute for Cognitive and Brain Health Northeastern University.

## Abstract

Understanding speech requires listeners to integrate incoming input with prior linguistic and thematic knowledge to access meaning, a task greatly aided by prediction. Surprisal and related phenomena (e.g., next word prediction) tend to be associated with broad activation of language regions during listening. A major challenge for interpreting the neural correlates of surprisal is that lexical expectancies are constrained both by syntactic and semantic information. In contrast, semantic distance yields more of a pure metric of relatedness between one language constituent (e.g., word, phrase, n-gram) and another within a high-dimensional semantic space. We contrasted surprisal and semantic distance in running discourse during naturalistic audiovisual language comprehension. We analyzed fMRI scans from 20 English-speaking adults who viewed the full-length film *500 Days of Summer*. Our focus was on brain sensitivity to either surprisal or semantic distance as language unfolded word-by-word. To this end, we implemented a series of hierarchical voxelwise encoding models. After accounting for low-level auditory and visual features, the addition of either surprisal or semantic distance improved prediction in bilateral superior temporal gyrus (STG), left inferior frontal gyrus, and right cerebellum. Critically, both semantic distance and surprisal explained unique variation in bilateral temporal cortex after accounting for the other. These results support the notion that the brain tracks both broad probabilistic prediction as well as semantic integration. More generally, these findings extend previous work on auditory-only speech by demonstrating that surprisal and semantic distance predict meaningful neural variation even within a rich audiovisual context.

## Introduction

Comprehending natural language requires listeners to integrate incoming acoustic input with prior linguistic and thematic knowledge to access meaning. These processes are supported by prediction, such that listeners continuously generate probabilistic expectations about upcoming information based on prior context (Brothers et al., 2023; Hale, 2001; Levy, 2008). Within this framework, the predictive difficulty of any given word can be operationalized as surprisal. While there are several methods that can be used to calculate surprisal, the most common definition is the negative logarithm of the probability of a word given its prior context (Hale, 2001). That is, upcoming words that are less predictable given their prior context are higher in surprisal.

Converging evidence across behavioral and neurophysiological measures supports the notion that the brain tracks the probabilistic structure of language as indexed by surprisal. Higher surprisal is associated with longer reading times (Levy, 2008; N. J. Smith & Levy, 2013), larger N400 amplitudes (Frank et al., 2015), and increased activity in bilateral temporal language regions (Brodbeck et al., 2018; Willems et al., 2016). More recently, large language models (LLMs) have been used to generate surprisal estimates (Goldstein et al., 2022; Michaelov et al., 2024; Russo et al., 2022). Unlike n-gram models, LLMs estimate probabilities based on learned transformer representations of prior context. LLM-derived surprisal estimates closely correlate with human predictability judgments (Szewczyk & Federmeier, 2022), and are associated with activation in the language network (Tuckute et al., 2024).

Despite wide use and intuitive appeal, surprisal can be difficult to interpret. It reflects a composite measure of predictability that blends multiple forms of linguistic constraint including word frequency, syntactic structure, and semantic coherence. Thus, it is impossible to link surprisal effects with specific psycholinguistic factors. Semantic distance, however, selectively captures the semantic integration component of predictability (Landauer & Dumais, 1997; Reilly et al., 2023; Rips et al., 1973). Semantic distance quantifies dissimilarity between concepts using a high-dimensional representation of semantic space, often learned from word co-occurrence statistics (Landauer & Dumais, 1997; Reilly et al., 2023, 2025). Using semantic distance to index conceptual similarity between a current word and its context in a continuous listening task, Mechtenberg and colleagues (2026) demonstrated that changes in semantic distance was associated with activation in the bilateral temporal gyrus and left inferior frontal gyrus.

Although semantic distance and surprisal are distinct constructs, both factors tend to be correlated and drive similar patterns of neural activation, most notably within the superior temporal gyrus (Giglio et al., 2026; Mechtenberg et al., 2026; Tuckute et al., 2024; Willems et al., 2016; cf., Frank & Willems, 2017). These findings raise the question of whether surprisal and semantic distance reflect a shared underlying neural representation within the STG, or whether the neural signatures to surprisal and semantic distance are dissociable. Dissociable effects between these measures may reflect distinct sensitivities to different levels of linguistic information, while shared effects may indicate that the surprisal-related response is largely driven by semantics.

The major focus of the present study was to contrast neural responses to semantic distance and surprisal. Additionally, to our knowledge, all previous work on surprisal and semantic distance has been conducted on auditory-only speech, which is not representative of real-world listening conditions. It remains unknown whether previous findings generalize to speech that is heard in a rich naturalistic audiovisual context with supporting information from the visual scene, which may provide additional constraints.

To address these questions, we examined how surprisal and semantic distance relate to neural responses while participants watched a full feature-length film during continuous functional MRI scanning (Aliko et al., 2020). Using a series of hierarchical voxelwise encoding models, we tested whether surprisal and semantic distance make unique contributions toward predicting neural responses over and above visual, auditory, and word-level features. We hypothesized that both surprisal and semantic distance would drive neural responses in the bilateral STG, while also explaining unique variance in neural activity, indicating that these metrics are complementary but partially dissociable.

## Method

### Participants

Data were taken from the Naturalistic Neuroimaging Database (NNDB; Aliko et al., 2020). Participants were 20 adults aged 19-53 (mean age = 27.70, SD = 10.14; 10 female, 10 male). All participants were native speakers of English, right-handed, without hearing impairment, with normal or corrected-to-normal vision, and with no history of neurological or psychiatric illness. All participants provided informed consent in accordance with institutional review board guidelines. None of the participants had previously seen the movie *500 Days of Summer*.

### Stimulus and Procedure

All participants viewed the movie *500 Days of Summer* (2009; duration 91.17 minutes) during continuous fMRI acquisition. Audio was routed through MR-compatible insert earphones. The film was presented in two 40-50 minute segments, with additional breaks if requested by the participant. For breaks, playback was paused and resumed with precise synchronization to the scanner trigger pulses. Participants were instructed to remain as still as possible and attend to the movie.

### MRI data Acquisition and Preprocessing

Imaging data were acquired using a 1.5 T Siemens MAGNETOM Avanto scanner using a 32 channel head coil. Functional images were obtained using a multiband echo-planar imaging (EPI) sequence (TR = 1 s, TE = 54.8 ms, flip angle = 75 degrees, 40 interleaved slices, 3.2 mm voxel size). T1-weighted anatomical images were acquired using a 10-minute MPRAGE sequence (TR = 2.73s, TE = 3.57 ms, 1 mm resolution).

Preprocessing was performed using AFNI and FSL tools (Cox & Hyde, 1997; Jenkinson et al., 2012). Functional data were corrected for slice timing differences (3dTshift), despiked (3dDespike), and motion corrected by aligning each volume to a reference (3dvolreg). Functional images were coregistered to each participant’s anatomical image and normalized to MNI space (3dNwarpApply). Spatial smoothing was applied to a target of 6 mm (3dBlurToFWHM). Time series were detrended and cleaned using a nuisance regression (3dTproject) which included motion parameters, white matter signal, cerebrospinal fluid signal, and low frequency drift. Independent component analysis was used to identify additional artifacts that were regressed out of the data (FSL MELODIC version 3.14). The time series were also corrected for delays introduced by breaks using interpolation-based shifting (Aliko et al., 2020). For more information about data acquisition and preprocessing, see Aliko and colleagues, 2020.

### Feature Extraction

#### Semantic Distance

We followed the procedures of Mechtenberg and colleagues (Mechtenberg et al., 2026), by using the SemanticDistance R package (Version 0.1.1 https://CRAN.R-project.org/package=SemanticDistance) to calculate semantic distance values for each word and to clean and format the transcript (Reilly et al., 2023). The movie transcript was cleaned by converting all words to lowercase, removing non-alphabetic characters, expanding contractions, removing proper nouns, and eliminating stopwords based on a custom stopword list (e.g., determiners, pronouns, prepositions). The SemanticDistance package uses an internal database of multidimensional semantic vectors for over 70,000 English words, which was created by training the GloVe semantic model (Pennington et al., 2014) on the Corpus of Contemporary American English (Davies, 2010). SemanticDistance functions were used to calculate semantic similarity between words, as measured by the pairwise cosine distance between vectors, with semantic distance calculated as 1 minus cosine similarity. In the present study, a rolling n-gram window of five words was used to determine the semantic distance between each word and the previous five words. This window size was chosen because of previous work which found little additional benefit to a larger window size beyond five content words (Mechtenberg et al., 2026). A portion of the time-series for semantic distance, as well as a visualization of other extracted movie features is shown in **Figure 1**.

**Figure 1.**
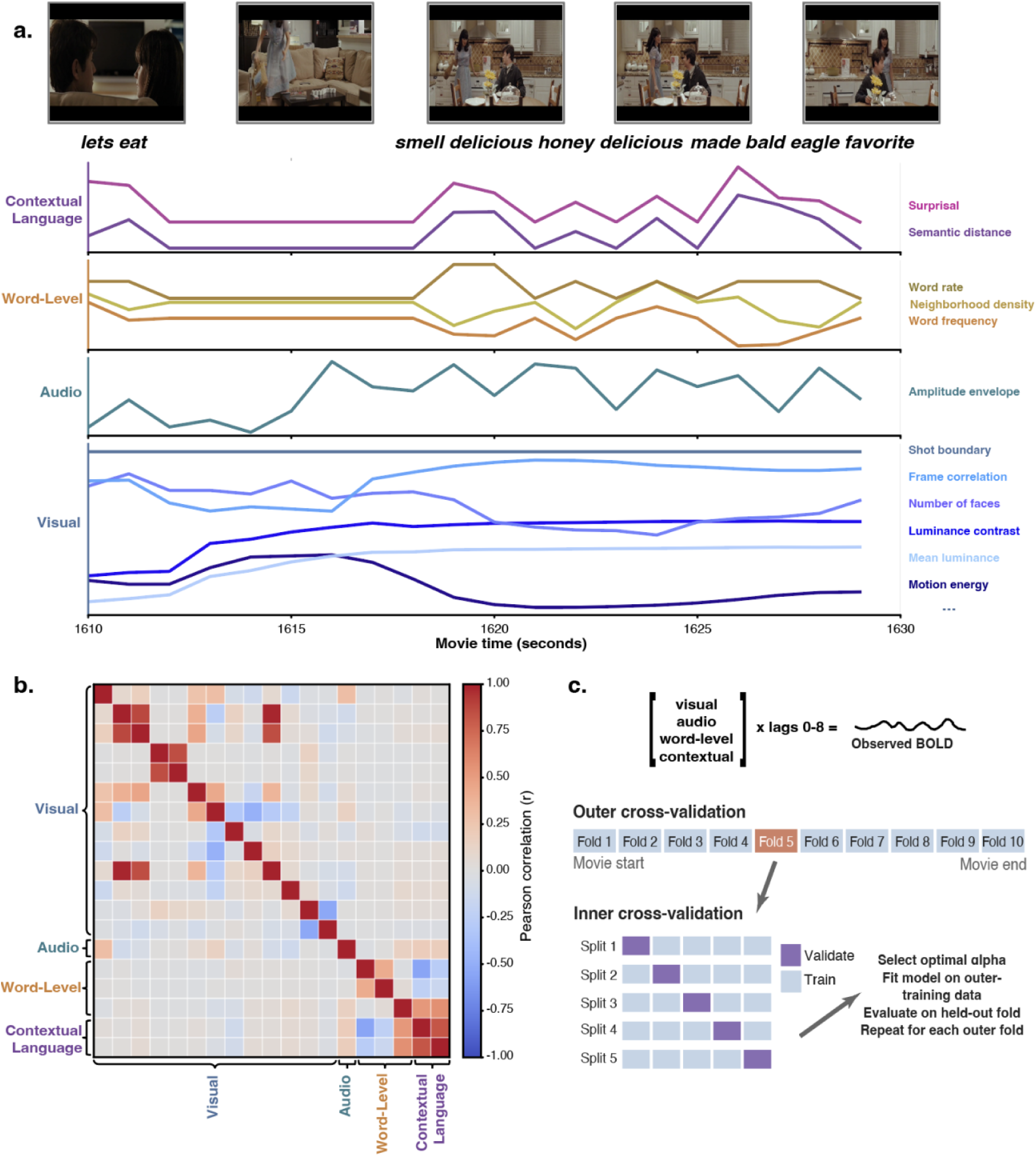
Feature extraction and modeling schematic. **a.** A sample of movie frames and aligned words from the movie transcript are shown for a 20 s segment of *500 Days of Summer*. Time-series for predictors downsampled to 1 s TRs are shown for the four feature sets of contextual language features, word-level features, the auditory amplitude envelope, and a sample of 6 of the visual features included in our model. These temporally aligned features were expanded with a finite impulse response filter and used in the hierarchical encoding models. **b.** Correlation matrix of stimulus features once downsampled to TR. **c.** Schematic of the nested cross-validation procedure used for each model. Features were expanded using lags of 0-8 TRs, and the movie was split into 10 outer folds. On each iteration one fold was held out as the test set while 5-fold cross-validation within the remaining training data was used to select the ridge regularization parameter. The model was then refit using all data in the outer training set and applied to the held-out fold. This procedure was repeated until each outer fold had been used as the test set.

#### Surprisal

Before the calculation of semantic distance, word-level surprisal was estimated using the pretrained GPT-2 language model (117M parameters) using the HuggingFace Transformers Library (Wolf et al., 2020). We note that while newer language models exist, newer models seem to generalize worse to human behavioral and neural data (Lin & Schuler, 2026; B.-D. Oh & Schuler, 2023). For each word in the movie transcript, we computed the conditional probability of that word given the preceding 10 words of context. The window size of 10 words was used to roughly equate the context window used to calculate surprisal with the 5-word window of semantic distance, as surprisal considers every word in the transcript and semantic distance only content words. The transcript was tokenized using GPT-2’s byte-pair encoding tokenizer. As words may consist of many tokens, word probability was calculated as the product of the sequential probabilities of each token within a word, conditioned on the preceding context. Surprisal was thus calculated as Surprisal(w*_i_*) = -log2*P*(w*_i_* | w*_i-10_*,…,w*_i-1_*), where w*_i_* is the current word in the transcript.

#### Visual, Auditory, and Word-level Features

A series of visual control features was calculated using the OpenCV Version 4.6.0 computer vision library (Bradski & Kaehler, 2000): overall luminance, contrast, color properties in HSV space, edge and texture statistics (Sobel edge magnitude and Laplacian variance), optical-flow estimations of motion energy, and the detection of shot boundaries. First derivatives of luminance and contrast were included to index rapid visual changes. Additional regressors were included for higher-level visual confounds such as camera motion, as well as the presence and number of faces. The auditory amplitude envelope was included as a low-level predictor of auditory drive, calculated as short-time RMS amplitude using 25 ms windows with a 10 ms step size. Word-level controls included lexical frequency values obtained from the SUBTLEX-US database (Brysbaert & New, 2009), phonological neighborhood density defined as the number of phonological neighbors obtained from the English Lexicon Project (Balota et al., 2007), and word rate, defined as the number of content words occurring within each TR. All features were averaged within 1 s TR-linked epochs.

### Hierarchical Encoding Models

A series of voxelwise ridge regression encoding models were created to determine the contribution of the visual, auditory, and language features to the BOLD response. All predictors were expanded using a Finite Impulse Response approach with lags from 0– 8 TRs, to allow the model weights to capture the delayed hemodynamic response without assuming a canonical hemodynamic response function (Antonello et al., 2023). Our choice of lags is consistent with the suggestion that extending FIR windows beyond approximately 8 seconds yields diminishing or negative returns (Binhuraib et al., 2025).

A schematic of the model estimation procedure is shown in **Figure 1**. Model performance was evaluated using 10-fold cross-validation, such that the original 5470 TRs were separated into 10 blocks of 547 TRs each. To reduce temporal leakage due to autocorrelation and lagged predictors, training samples within 10 TRs of the boundary marking each held-out test block were excluded (Binhuraib et al., 2025; see Oota et al., 2023). Within each fold, both predictors and voxel time series were standardized using only parameters estimated from the training data. Hyperparameter selection (ridge regularization alpha) was determined using a nested inner 5-fold blocked cross-validation, with alpha selected from a logarithmically spaced grid ranging from 10^−2^ to 10^9^. For each voxel, model performance was quantified as the cross-validated correlation between observed and predicted BOLD responses (e.g., Huth et al., 2016). Incremental effects were computed by subtracting prediction performance using hierarchically nested models.

We used a hierarchical encoding procedure to determine the incremental contribution across feature sets. The baseline model, Model 1, included all of the visual features extracted using OpenCV. Model 2 included all of the features of Model 1 as well as the acoustic amplitude envelope. Model 3 contained all of the features of Model 2 as well as the word-level features of word rate, lexical frequency, and phonological neighborhood density. Model 4 contained all of the features present within Model 3, as well as the predictive language features of semantic distance and surprisal. To examine shared and unique contributions of surprisal and semantic distance, two incremental models were created between Model 3 and Model 4, which excluded either semantic distance or surprisal. Thus, the unique effect of semantic distance was estimated as the performance of the full model minus the model containing surprisal but not semantic distance, and the unique effect of surprisal was calculated as the performance of the full model minus the model containing semantic distance but not surprisal.

Group-level nonparametric inference was used to test whether prediction performance or incremental model improvements reliably differed from zero across participants. For each map, we used a one-sample test against zero using 10,000 sign-flip permutations and threshold free cluster enhancement (TFCE; (Abraham et al., 2014; Nichols & Holmes, 2002; S. M. Smith & Nichols, 2009)). Statistical significance was assessed using family-wise error correction, with significance defined as TFCE-corrected *p* < .05. For visualization, group mean maps were masked by the TFCE-corrected significance map, such that only voxels surviving correction were displayed. Group-level volumetric maps were projected to the fsaverage cortical surface using Nilearn (Abraham et al., 2014) and to a cerebellar flatmap using SUIT (Diedrichsen, 2006; Diedrichsen & Zotow, 2015; Wang et al., 2026).

## Results

We first examined the spatial distribution of prediction performance across the four hierarchical encoding models to see how well the increasingly linguistically relevant stimulus feature sets predict neural responses during movie watching. This analysis was conducted to determine whether the model hierarchy produced the expected visual, auditory, and language-related patterns of predictivity, and also to test whether the addition of semantic distance and surprisal improved model performance even after accounting for lower level visual, auditory, and word-level features.

**Figure 2** displays the results of the hierarchical encoding modeling procedure, displaying both the full spatial distribution of predictivity, as well as areas where a statistically significant change in predictivity from the prior model was detected. Model 1, which contained visual features (framewise luminance, RMS contrast, HSV color statistics, edge magnitude, Laplacian variance, optical flow magnitude, shot boundary estimates, face count, and temporal derivatives of luminance and contrast), predicted broad posterior neural activity, particularly in occipital regions including primary visual cortex. One large cluster spanning 11202 voxels was identified, with peaks in the right and left occipital pole, left intracalcarine cortex, and the superior division of the right lateral occipital cortex.

**Figure 2.**
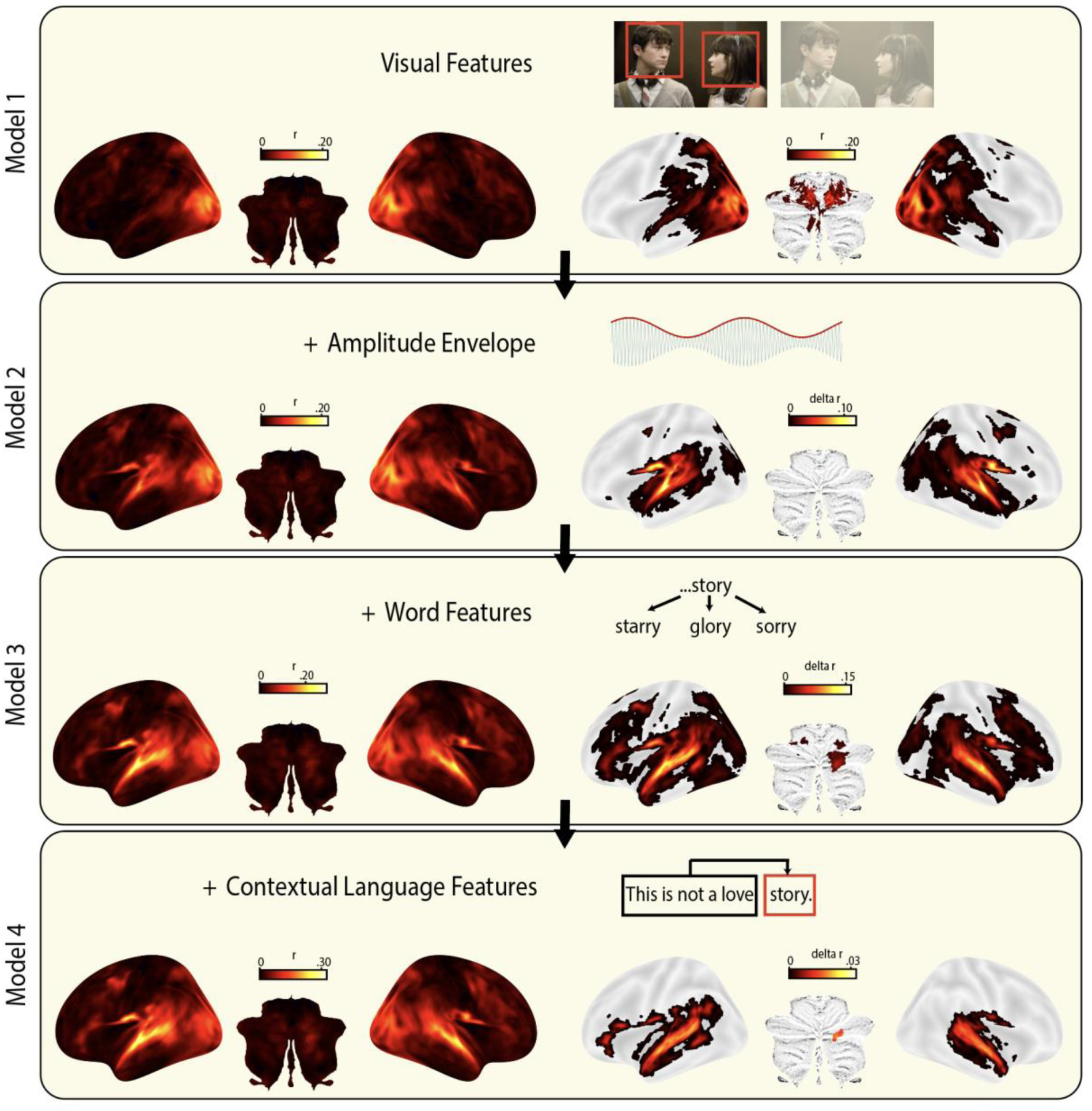
Hierarchical Modeling Results. Results from the hierarchical encoding modeling procedure are shown for Models 1-4. Within each panel, the left displays uncorrected predictivity for the overall model plotted on a cortical surface, the top displays a visual example of each regressor class, and the right displays the change in predictivity from the previous modeling step including only voxels where a statistically significant change was detected (FWE corrected TFCE *p* < 0.05). Note that for Model 1, as there is no previous model to compute change, any predictivity that was statistically significant is displayed.

The addition of the auditory amplitude envelope in Model 2 extended model predictivity to bilateral temporal regions, with a significant increase in model performance, visually notable within bilateral STG. There were 19 clusters identified as associated with increased predictivity due to the addition of the auditory amplitude envelope. The largest two clusters were in the left and right temporal cortex, with peaks in the planum temporale, planum polare, and STG.

Model 3 introduced three word-level features: word frequency, phonological neighborhood density, and content word rate. The addition of these three features resulted in statistically significant increases in predictivity throughout broad portions of the occipital, frontal, and temporal lobes. Nine clusters were identified, with the largest clusters containing peaks in left and right temporal lobe, including both anterior and posterior divisions of the STG. A peak was also identified in the cerebellum at right crus II, in bilateral precuneus, and bilateral superior frontal gyrus.

Of primary interest, Model 4 contained all of the features present in previous models, and also included the two context-sensitive language features (semantic distance and surprisal). While the addition of context-sensitive measures largely did not change the spatial extent of predictivity, this model achieved the highest performance, particularly in the temporal cortex. The regions in which model performance increased significantly are shown in **Table 1**, which comprised four clusters. The largest cluster appeared in the left temporal cortex, with a peak in the anterior division of left STG, and with subpeaks in the posterior division of the STG and the left temporal pole. The second largest cluster appeared in the right temporal cortex, with a peak in the anterior division of the right STG with subpeaks in the posterior division and right temporal pole. A cluster also emerged within the cerebellum with a peak in right crus II, and a fourth, smaller cluster in left Heschl’s gyrus.

**Table 1.** Peak change in predictive performance when both semantic distance and surprisal were added to the model.

| Region | Cluster size (voxels) | Cluster size (mm<sup>3</sup>/uL) | Peak Value (delta r) | x | y | z |
| --- | --- | --- | --- | --- | --- | --- |
| L Superior Temporal Gyrus, anterior division | 1185 | 31995.0 | 0.02521 | -61.5 | 1.5 | -7.5 |
| L Superior Temporal Gyrus, posterior division |  |  | 0.02484 | -67.5 | -28.5 | 1.5 |
| L Supramarginal Gyrus, posterior division |  |  | 0.0237 | -64.5 | -43.5 | 7.5 |
| L Superior Temporal Gyrus, posterior division |  |  | 0.02273 | -55.5 | -40.5 | 4.5 |
| L Superior Temporal Gyrus, posterior division |  |  | 0.02151 | -67.5 | -31.5 | 10.5 |
| L Temporal Pole |  |  | 0.01782 | -49.5 | 22.5 | -19.5 |
| R Superior Temporal Gyrus, anterior division | 821 | 22167.0 | 0.02621 | 58.5 | -4.5 | -7.5 |
| R Superior Temporal Gyrus, posterior division |  |  | 0.02491 | 61.5 | -13.5 | -4.5 |
| R Superior Temporal Gyrus, posterior division |  |  | 0.02462 | 61.5 | -25.5 | 1.5 |
| R Temporal Pole |  |  | 0.01973 | 58.5 | 7.5 | -10.5 |
| R Temporal Pole |  |  | 0.01795 | 49.5 | 16.5 | -25.5 |
| R Superior Temporal Gyrus, posterior division |  |  | 0.01627 | 49.5 | -13.5 | -10.5 |
| Right Crus II | 26 | 702.0 | 0.01909 | 25.5 | -88.5 | -31.5 |
| L Heschl's Gyrus (includes H1 and H2) | 5 | 135.0 | 0.00889 | -40.5 | -22.5 | 10.5 |
| L Insular Cortex |  |  | 0.0062 | -34.5 | -25.5 | 16.5 |

Note. Only voxels surviving TFCE-based familywise-error correction are included. Clusters represent 26-connected components of significant voxels and subpeaks were required to be at least 8 mm apart. Coordinates are reported in MNI space.

As previously mentioned, two additional models were constructed for the purpose of determining the incremental and unique contributions of surprisal and semantic distance via model comparisons. The results of this procedure are shown in **Figure 3**. We found that when adding either semantic distance or surprisal to Model 3, predictivity increased in similar regions, including the bilateral superior temporal cortex and left inferior frontal gyrus, suggesting that both measures captured language specific variance beyond the visual, auditory, and lexical controls. Clusters with peaks in the left and right STG, the left frontal gyrus, and the cerebellum at right crus II were identified in both maps (see supplemental material).

**Figure 3.**
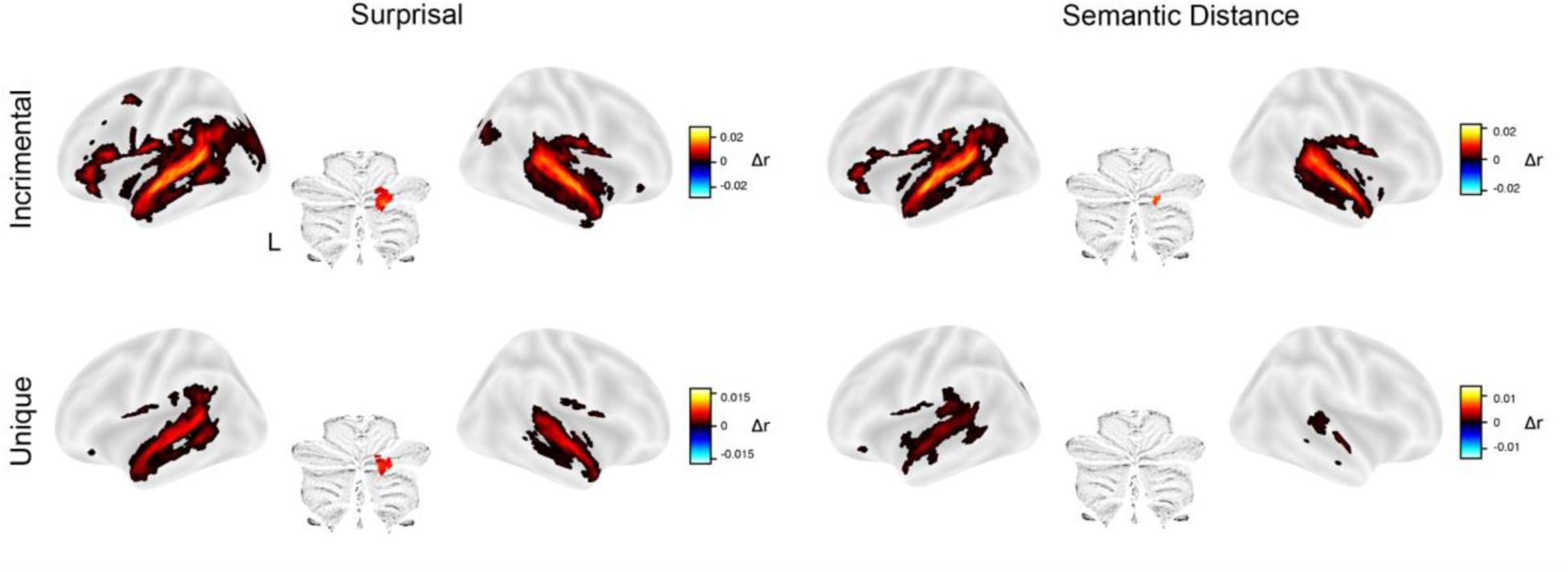
Incremental and unique predictivity for Surprisal and Semantic Distance. Results from the incremental addition of Semantic Distance compared to Model 3 is shown on the top left. Change in predictivity with the addition of Surprisal compared to Model 3 is shown on the top right. The bottom left panel shows unique contributions of semantic distance as compared to Model 3 + surprisal, while the bottom right panel shows the unique contribution of surprisal compared to Model 3 + semantic distance. For all images, voxels are displayed only where a statistically significant change was detected (FWE corrected TFCE *p* < 0.05).

However, when comparing these incremental models to Model 4 to assess unique contributions, some differences emerge, which is visualized in the lower half of **Figure 3**. While both predictors explain unique predictivity in the temporal lobe, surprisal increased predictivity in the right cerebellum over and above the contribution of semantic distance, and increased model predictivity more broadly in the right STG. The clusters which emerged from the unique contribution of semantic distance are shown in **Table 2**, with the largest clusters containing peaks identified in left STG (with two smaller clusters in the right STG). **Table 3** shows the clusters which were associated with the unique contribution of surprisal, with two wide clusters in the left and right temporal cortex, as well as smaller clusters in the cerebellum (right crus I and II).

**Table 2.** Peak change in predictive performance due to semantic distance over and above the contribution of surprisal.

| Region | Cluster size (voxels) | Cluster size (mm<sup>3</sup>/uL) | Peak Value (delta r) | x | y | z |
| --- | --- | --- | --- | --- | --- | --- |
| L Superior Temporal Gyrus, anterior division | 367 | 9909.0 | 0.00444 | -55.5 | -4.5 | -7.5 |
| L Parietal Operculum Cortex |  |  | 0.00395 | -58.5 | -28.5 | 13.5 |
| L Superior Temporal Gyrus, anterior division |  |  | 0.00378 | -61.5 | 1.5 | -1.5 |
| L Temporal Pole |  |  | 0.00373 | -52.5 | 19.5 | -19.5 |
| L Planum Temporale |  |  | 0.00364 | -55.5 | -19.5 | 4.5 |
| L Superior Temporal Gyrus, posterior division |  |  | 0.00358 | -61.5 | -25.5 | 1.5 |
| R Superior Temporal Gyrus, posterior division | 23 | 621.0 | 0.00338 | 61.5 | -28.5 | 4.5 |
| L Parietal Operculum Cortex | 22 | 594.0 | 0.00398 | -43.5 | -37.5 | 19.5 |
| L Planum Temporale |  |  | 0.00358 | -40.5 | -34.5 | 10.5 |
| R Superior Temporal Gyrus, anterior division | 17 | 459.0 | 0.00334 | 64.5 | -4.5 | -4.5 |
| R Superior Temporal Gyrus, posterior division |  |  | 0.00315 | 67.5 | -10.5 | 1.5 |
| L Lateral Occipital Cortex, superior division | 8 | 216.0 | -0.00345 | -16.5 | -88.5 | 28.5 |

Note. Only voxels surviving TFCE-based familywise-error correction are included. Clusters represent 26-connected components of significant voxels and subpeaks were required to be at least 8 mm apart. Coordinates are reported in MNI space.

**Table 3.** Peak change in predictive performance due to surprisal over and above the contribution of semantic distance.

| Region | Cluster size (voxels) | Cluster size (mm<sup>3</sup>/uL) | Peak Value (delta r) | x | y | z |
| --- | --- | --- | --- | --- | --- | --- |
| L Middle Temporal Gyrus, temporooccipital part | 719 | 19413.0 | 0.00898 | -64.5 | -43.5 | 4.5 |
| L Superior Temporal Gyrus, posterior division |  |  | 0.00827 | -64.5 | -28.5 | 1.5 |
| L Superior Temporal Gyrus, anterior division |  |  | 0.00796 | -61.5 | 1.5 | -7.5 |
| L Superior Temporal Gyrus, posterior division |  |  | 0.00796 | -58.5 | -22.5 | -1.5 |
| L Superior Temporal Gyrus, posterior division |  |  | 0.00747 | -55.5 | -40.5 | 4.5 |
| L Superior Temporal Gyrus, posterior division |  |  | 0.00707 | -64.5 | -16.5 | 1.5 |
| R Superior Temporal Gyrus, anterior division | 509 | 13743.0 | 0.00838 | 61.5 | -4.5 | -4.5 |
| R Superior Temporal Gyrus, posterior division |  |  | 0.00809 | 58.5 | -13.5 | -4.5 |
| R Superior Temporal Gyrus, posterior division |  |  | 0.0074 | 61.5 | -25.5 | 1.5 |
| R Superior Temporal Gyrus, posterior division |  |  | 0.00687 | 70.5 | -13.5 | 1.5 |
| R Superior Temporal Gyrus, posterior division |  |  | 0.00673 | 70.5 | -19.5 | 7.5 |
| R Temporal Pole |  |  | 0.00646 | 49.5 | 16.5 | -25.5 |
| Right Crus II | 33 | 891.0 | 0.00801 | 19.5 | -82.5 | -34.5 |
| Right Crus I | 2 | 54.0 | 0.00507 | 16.5 | -79.5 | -22.5 |

Note. Only voxels surviving TFCE-based familywise-error correction are included. Clusters represent 26-connected components of significant voxels and subpeaks were required to be at least 8 mm apart. Coordinates are reported in MNI space.

## Discussion

Understanding natural language requires the listener to continuously stitch incoming words with previously heard linguistic context. In the present study, we tested whether surprisal and semantic distance, two computational measures of contextual language processing, explain distinct neural responses during naturalistic audiovisual comprehension. This was accomplished by using voxelwise encoding models on fMRI data collected while participants viewed a full-length feature film. We found that adding context-sensitive linguistic features improved prediction of neural responses beyond modeled visual, auditory, and word-level stimulus features. Of special focus, both surprisal and semantic distance explained neural variation after accounting for the other, suggesting that these features are non-redundant.

The systematic improvement of model performance across hierarchical models provides an internal validation of our modeling approach. In brief, we found that adding increasingly linguistically relevant features to our model predicted progressively broader portions of cortex. More specifically, visual features primarily predicted activity in posterior visual regions, the addition of the auditory amplitude envelope increased predictivity in bilateral STG, and word-level features improved prediction throughout cortex, particularly in temporal and frontal language regions. Adding surprisal and semantic distance improved prediction beyond these visual, auditory, and word level controls, predominantly in the bilateral temporal regions and left inferior frontal gyrus. This progression suggests that the impact of surprisal and semantic distance on the prediction of neural responses cannot be explained by variation in broad visual input, acoustic amplitude, word rates, or lexical features.

The surprisal findings converge with evidence from several studies which demonstrate that neural responses during language comprehension are sensitive to the probability of incoming words given their prior linguistic context (Brodbeck et al., 2022; Frank et al., 2015; Tuckute et al., 2024; Willems et al., 2016). Using EEG, Frank et al. (2015) demonstrated that word surprisal correlated with N400 amplitude during sentence reading. In a naturalistic listening fMRI paradigm, Willems et al. (2016) found activity across temporal and inferior frontal regions, while Brodbeck et al. (2022) showed responses to lexical surprisal based on both local and broader sentence context in bilateral temporal lobe using MEG. More recently, Tuckute et al. (2024) demonstrated that global sentence-level surprisal was a primary determinant of language-network responses. The present study replicated the effects of surprisal in bilateral STG and left inferior frontal gyrus, and demonstrated that this response remains robust even in audiovisual contexts.

Semantic distance has received relatively less attention than surprisal in imaging research, particularly with fMRI. Electrophysiological studies have shown that semantic dissimilarity predicts responses similar to an N400 during continuous narrative listening (Broderick et al., 2018), and that semantic similarity can enhance early encoding of acoustic and phonological features (Broderick et al., 2019). In fMRI, semantic distance during a podcast listening task was associated with a broad bilateral frontotemporal network, including bilateral superior and middle temporal cortex, left inferior frontal gyrus, and bilateral cerebellum (Mechtenberg et al., 2026). The present results replicate the findings of Mechtenberg and colleagues (2026) using a naturalistic movie watching paradigm, and demonstrate that model prediction performance on held out data can be improved by the incorporation of semantic distance over and above visual, auditory, and word-level features.

Of primary interest was whether surprisal and semantic distance make statistically unique contributions to neural predictivity. We found that, despite producing effects in regions of cortex that were largely overlapping, both measures accounted for unique variation in bilateral STG. A plausible interpretation is that semantic distance and surprisal influence processing at multiple stages of the speech-language processing hierarchy (Brodbeck et al., 2022; Broderick et al., 2019). Bilateral STG is often associated with acoustic-phonetic analyses that support recognition of word forms (Davis & Johnsrude, 2003; Hickok & Poeppel, 2007), and the unique STG contributions of semantic distance and surprisal may reflect the use of context to facilitate the processing of word-form details through top-down feedback. Consistent with this interpretation are fMRI findings that prior expectations can modulate speech responses in superior temporal cortex (Blank & Davis, 2016). MEG results also indicate that prior evidence can rapidly influence cortical representations of spectrotemporal speech information (Sohoglu & Davis, 2020), potentially reflecting top down influences from the inferior frontal gyrus on speech representations within the STG (Gagnepain et al., 2012; Sohoglu et al., 2012).

We also found a region of right cerebellum (Crus I and II) that was associated with surprisal. While unexpected, this finding is consistent with the notion that the cerebellum supports linguistic prediction and processing of prediction error during language comprehension (Lesage et al., 2017; Mechtenberg et al., 2024; Moberget et al., 2014). It is also part of an increasing awareness of the wide variety of representations in the cerebellum, related to lexical processing (Mechtenberg et al., 2024), semantic memory (LeBel et al., 2021), and naturalistic stimuli (King et al., 2019).

The present findings both converge with and depart from previous studies that also modeled surprisal and semantic distance simultaneously in language comprehension. Frank and Willems (2017) found that semantic distance and surprisal explained neural variation after accounting for the other. However, the effects observed in their study were more spatially dissociable, as they found that surprisal was associated with activation in bilateral STG and left posterior fusiform gyrus while semantic distance was associated with left anterior temporal cortex, bilateral angular gyrus, and precuneus. Relatedly, Russo and colleagues (2020) compared lexical surprisal with a semantics-weighted surprisal measure during narrative listening, and found that the model which incorporated semantic similarity into the calculation of word surprisal provided better fit than lexical surprisal in left superior and middle temporal cortex, right anterior temporal cortex, and left inferior frontal gyrus. However, because semantic similarity and lexical surprisal were entered separately into models rather than entered simultaneously as independent predictors, their analysis could not determine whether semantic distance itself explained neural responses over and above surprisal.

Methodological choices may have contributed to differences between the present study and prior studies. The three studies had different operationalizations of contextual linguistic information: the present study used transformer-based GPT-2 surprisal and GloVe-based semantic distance, Russo and colleagues (2020) used a trigram model for lexical surprisal that was scaled by a semantic similarity factor, and Frank and Willems (2017) used an n-gram and skip-bigram model with semantic distance estimated from skip-gram word embeddings. The three studies also applied different statistical procedures, varied in what regressors were used for control, and varied in modality, as both prior studies were conducted in a unimodal context. Relative to earlier approaches, the present study provides a stricter test of the independent contributions of these linguistic features by using a more context sensitive estimate of surprisal and evaluating incremental out-of-sample neural prediction.

It is notable that effects of semantic distance in both this study and Mechtenberg and colleagues (2026) were not seen in the broader network of regions typically implicated in semantic processing. Studies of semantic processing using words and pictures have been associated with activation in anterior and middle temporal cortex, angular and inferior parietal cortex, ventral temporal cortex, inferior frontal gyrus, the posterior cingulate and medial prefrontal cortex (Binder et al., 2009; Lambon Ralph et al., 2017; Patterson et al., 2007; Vandenberghe et al., 1996). Distributed semantic neural representations for words have also been observed using movie stimuli (e.g., Huth et al., 2016). We suggest, however, that semantic distance and surprisal do not index the representation of conceptual knowledge *per se*, but the context-dependent integration of words. That is, while the neural representation of semantic information may be distributed across a wide cortical network, the consequences of that information for ongoing speech comprehension are expressed most strongly as a modulation of acoustic-phonetic and lexical representations within the STG.

There are several limitations to the present study. This study was limited to one romantic comedy movie, and results should not be assumed to generalize across all genres (Westfall et al., 2016; Yarkoni, 2020). We also note that surprisal and semantic distance were highly correlated, and observed naturally as they occurred in the movie rather than experimentally manipulated. Future work could manipulate these features independently while controlling the visual input to determine whether their statistically unique predictive effects truly reflect dissociable neural computations rather than differences in how the features were operationalized or covariance with unmodeled features within the movie stimulus.

In conclusion, we examined the neural tracking of contextual language features during naturalistic movie viewing using a series of voxelwise hierarchical encoding models. We found that semantic distance and surprisal improved prediction performance beyond visual, auditory, and word-level stimulus features. Although the neural effects of the two measures were largely overlapping in anatomical space, both measures accounted for unique variation. These results indicate that both probabilistic and semantic relationships with prior heard content provide complementary but non-redundant information about neural activity during language processing, which remain relevant even in the presence of complex multimodal input.

## Acknowledgements

Work reported here was funded in part by grants R01 DC019507 and R01 DC013063 from the US National Institutes of Health.

